# Reconstructing creative thoughts: Hopfield Neural Networks

**DOI:** 10.1101/2023.09.18.557700

**Authors:** Denisa Checiu, Mathias Bode, Radwa Khalil

**Affiliations:** School of Electrical and Computer Engineering, Constructor University, Bremen, Germany; School of Business, Social, and Decision Sciences, Constructor University, Bremen, Germany

**Keywords:** Creative Thinking, Hopfield Neural Network, Patterns, Memory, Associative Chains

## Abstract

From a brain processing perspective, the perception of creative thinking is rooted in the underlying cognitive process, which facilitates exploring and cultivating novel avenues and problem-solving strategies. However, it is challenging to emulate the intricate complexity of how the human brain presents a novel way to uncover unique solutions. One potential approach to mitigate this complexity is incorporating creative cognition within the evolving artificial intelligence systems and associated neural models. Hopfield Neural Networks (HNN) are commonly acknowledged as a simplified neural model, renowned for their biological plausibility to store and retrieve information, specifically patterns of neurons. Therefore, we propose for the first time using HNN to generate a potential model that emulates specific cognitive processes of creative thinking based on generating meaningful links between seemingly disparate concepts.

## Introduction

Recent years have witnessed substantial advances and tremendous growth in artificial intelligence (AI) systems and their applications (Kappas & Gratch, 2023; Virvou, 2022; Zhang et al., 2022). The groundbreaking progress of conversational AI such as ChatGPT, which broke significant barriers of automation in communication (Abdullah et al., 2022; Wang et al., 2023), and the emerging era of human-AI interactions, in which our computers act as autonomous agents capable of making decisions (Kappas & Gratch, 2023; Virvou, 2022), is indisputable.

AI research can be classified into two primary categories: the emulation category, which seeks to understand human perception, cognition, and motor abilities (e.g., humanoid robots), and the application category, which seeks to use AI methods to model and create large-scale products (e.g., teleoperated devices)(Kappas & Gratch, 2023; Shneiderman, 2020).

This article focuses on emulation because we aimed to employ AI to study creative cognition. Through the creative act, we can perceive the world differently, find hidden patterns, and construct connections between seemingly unrelated events to generate extraordinary outputs (Abraham, 2018; Herrmann, 1981; Khalil & Demarin, 2023; Khalil & Moustafa, 2022). Mendelson’s (1974) model of knowledge access employed the concepts of defocused and focused attention to elucidate the semantic hierarchy of creative connections. The concept of the associative foundation of the creative process aligns with the earlier introduction of this notion by Mednick (1962).

It is widely accepted that creative products should express novel, valuable, and may be surprising ideas (Corazza, 2016; Gardner, 1994; Runco & Jaeger, 2012; Torrance, 1974; Wilson et al., 1954). Consequently, there is a broad consensus that creative thinking is recognized for its ability to generate solutions that are both novel, useful, and surprising (Han et al., 2021; Khalil & Moustafa, 2022; Mastria et al., 2022).

Nevertheless, creativity is multifaceted: it can be expressed in various forms, hues, and shapes and is, thus, enormously challenging to pin down (Abraham, 2013; Gaut, 2010; Kaufman & Beghetto, 2009; Sawyer, 2006, 2011; Squalli & Wilson, 2014; Tardif & Sternberg, 1988). One possibility to reduce this complexity and overlapping is to focus on the neural network architectures and their relevant learning rules, as Khalil and Moustafa (2022) suggested.

Therefore, we intend to incorporate this creative cognitive process-based semantic association (Beaty & Kenett, 2023; Kenett & Beaty, 2023; Li et al., 2021) into a conceptual framework for a creative, plausible neural network system using the Hopfield neural network (HNN). The biologically inspired mathematical tool HNN, developed by John J. Hopfield in 1982, is best recognized for its ability to store and recall knowledge (McEliece et al., 1987). Establishing the HNN model was remarkable in the neural computational and modeling era (Abe, 1989; Hopfield, 1982; KÖksal & Sivasundaram, 1993; Ramsauer et al., 2021).

HNN is a recurrent associative neural network model that is featured by having input from all neurons, and thus, it can retrieve memories from partial or noisy input (Hopfield, 1982). After receiving an input, the HNN uses iterative updating to update the neuron states synchronously or asynchronously to converge toward one of the stored memories and reach a stable state that recalls the information it was trained (Krotov & Hopfield, 2016). Therefore, HNN has been used for associative memory tasks, including associating input with its most similar memorized pattern or identifying unknown systems by computing the learning mechanism’s optimized coefficients to fit the nonlinear system (McEliece et al., 1987; Wang et al., 2003, 2005).

Using this network structure strongly resembles the architecture of human memory, which encodes several neurons that fire together in response to certain “microfeatures”(Gabora, 2010). In other words, the HNN recalls patterns by considering the correlation between the neurons using the weights matrix, not by storing the patterns; thus, we chose the HNN model to develop a semantic association-based creative conceptual framework (Beaty et al., 2023; Kenett & Beaty, 2023; Luchini et al., 2023).

We describe this semantic linkage as a radio knob that we can turn to tune whether the network can solve problems creatively or if it shuts down entirely; the threshold is used as a parameter. We refer to it as the “**first knob of creativity”**; in this context, we are invoking the “**second knob of creativity”** to facilitate exploring alternative solutions inside our network. By manipulating the knob, it is possible to selectively suppress specific patterns, facilitating the creative functioning of the HNN and identifying other patterns with which our input can be linked.

## Methods

This section describes the neural network structure and implementation of HNN (Fig.1).

**Figure 1.**
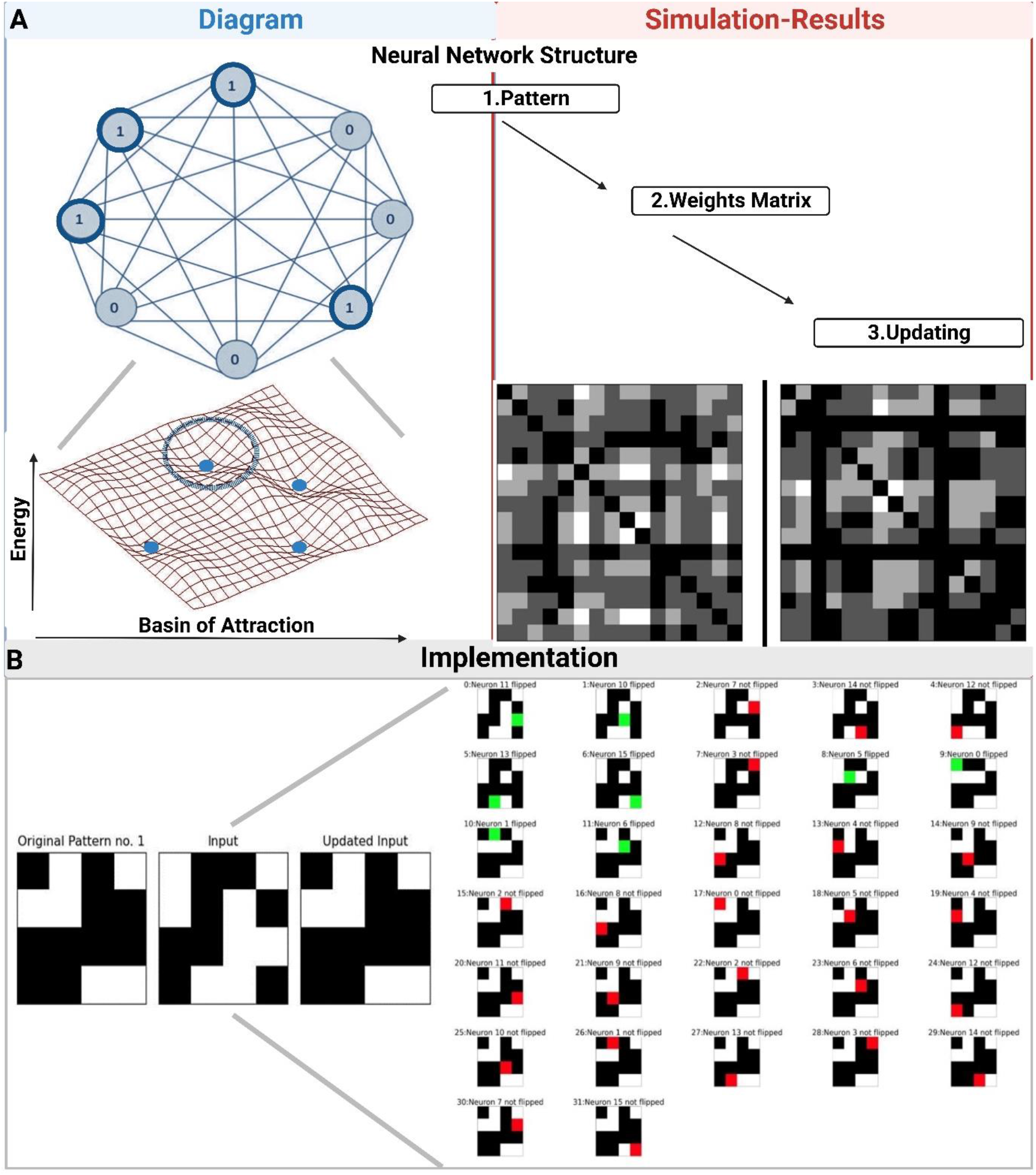
Description of the neural network structure of Hopfield neural networks (HNN) and its implementation. Panel A depicts a schematic representation of a binary HNN, illustrating its connectivity and the potential states of the neurons, which can be either on (active) or off (inactive). The lower graph depicted in panel A refers to the energy structure of HNN with multiple point attractors; fixed points are shown as small blue dots, and a dotted circle denotes a basin of attraction. The network consists of a pattern and a weight matrix [W], which directs its capacity to undergo single-step training as the computation is executed in a singular operation. The diagonal symmetry of W can be attributed to the symmetric and bi-directional connections among the neurons, which is a consequence of the complete connectivity. The lower graph of panel A on the right side illustrates visual representations of the weight matrix obtained from preliminary experiments conducted on binary HNN implementation. Panel B illustrates the graphical visualizations of the successful implementation of HNN. The graph on the left-hand side shows a randomly chosen pattern, its perturbation as the input fed into the HNN, and the output after two complete cycles of asynchronous updates from the initial experiments. The graph on the right-hand side represents asynchronous neuron updates of an input iterated through the network - a green cell means the neuron was updated, while a red cell translates to it staying the same value.

## Neural Network Structure

The network structure includes patterns and weight matrix updating (Fig.1a).

### 1. Patterns

We define n patterns, each length N; every pattern pi, with i = [1, n], is considered a memory network, which will learn during the training process and later reconstruct given an input close to or derived from it, essentially an input resembling it (Fig.1a).

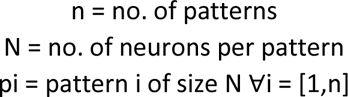

A pattern pi comprises N neurons, every neuron having a state, on or off. We used a binary HNN; a neuron can be either 0, meaning the neuron is not firing, or 1, the neuron is firing. This binary mode reflects a simplistic and discrete view of how the neurons act (Šíma et al., 2000). Therefore, for a general definition, we can consider each pattern as a random combination of 0s and 1s, each representing a neuron, forming a row vector.

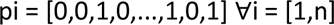

Each neuron is connected to all the others from the network, indicating that each connection has a unique weight, similar to the synapses between the neurons, representing communication between neurons and how these change their states accordingly (Ramsauer et al., 2021). An ensemble of specific neurons from different areas firing together in a unique sequence refers to a memorized concept (Hopfield, 1982).

We added another variable that can control the number of ones we have in all patterns - thus, we ensure that patterns are still randomly generated, but we know how many neurons are firing. We started by generating patterns with 10% of ones out of the total of N neurons (N = 1024). Given a noisy input, we were ready to perform a more thorough analysis of the range of thresholds for each n for which the network can adequately reconstruct patterns.

We performed 50 tests for each possible threshold value, ranging from 0 to 150 with increments of 10, for each number of patterns n, starting from 1 until 9. A test consisted of generating n new patterns, which contained 10% of randomly firing neurons, and then perturbing by 100 neurons a randomly chosen pattern, which we fed into the network as input.

Our goal is to develop a neurocomputational model for creative thinking based on semantic associations (Beaty et al., 2023; Benedek et al., 2023; Gabora, 2010; Gerver et al., 2023); it is logical to have less correlated concepts to identify the unexplored connections in between. Based on this reasoning, our patterns represent unrelated concepts; therefore, we do not require a large percentage of neurons firing per pattern (i.e., meaning less likelihood for correlation)(Amari & Maginu, 1988; Anishchenko & Treves, 2006; Hopfield, 1982; McEliece et al., 1987).

### 2. Weights Matrix

We structure the connectivity of the neurons as a matrix with defined weights. We included *n* patterns in our HNN to store and memorize; this process is performed through the computation of the weights matrix *W*.

We used the Hebbian learning rule to store the patterns in HNN. The fundamental concept underlying the Hebbian rule is encapsulated in the widely recognized phrase,**” Neurons that fire together wire together. Neurons that fire out of sync fail to link”**, meaning if two neurons fire simultaneously, their connectivity value - the unique weight connecting them - increases, and in return, causes their activity to become more correlated over time (Morris, 1999).

The reasoning above may be applied to the alternative scenario, wherein the neurons exhibit contrasting states. In this situation, their connection value diminishes and reaches a null state, specifically in our context. This tendency facilitates the convergence of inputs towards the most similar pattern, signifying the synchronized firing of neurons and indicating the connection between them (Amit, 1995; Brunel, 1996).

Based on this Hebbian rule, we created *W*; thus, the weights matrix is computed by summing up the outer product of each pattern *p_i_* - to measure the correlation between the patterns and our input.

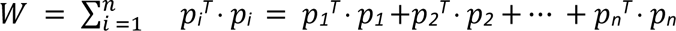

We considered each pattern *p_i_* a row vector, and as such *p^T^_i_* is a column vector, the outer product computation of a pattern resulting in a matrix.

One of the remarkable advantages of the HNN is its ability to train in one step, as the computation of the weights matrix *W* is completed in one operation (Abe, 1989; Abe & Gee, 1995; Atencia et al., 2005; Joya et al., 2002; KÖksal & Sivasundaram, 1993; Ramsauer et al., 2021). We used the diagonal symmetry of *W*, caused by the symmetric and bi-directional connections of the neurons, a direct result of the full connectivity (Fig.1a).

### 3. Updating

Given a binary input *x* of *N* neurons, we iterate it through HNN and make it converge towards a stable state. This convergence is facilitated by the utilization of a weights matrix *W*. Notably, the patterns themselves are not required for this operation, as the network has been trained, and all the necessary information is encapsulated within *W* (Ramsauer et al., 2021; Šíma et al., 2000).

There are two ways to perform an update: synchronously, meaning all of the neurons from the input *x* get updated in one iteration (Kobayashi, 2021; Michel et al., 1990), and asynchronously, where at each iteration, a random neuron (out of the *N* available ones) is chosen and updated (Aviel et al., 2004; Hopfield, 1982; Ramsauer et al., 2021). We used asynchronous updates to comprehensively examine the intermediate steps in reaching a solution. Consequently, an iteration is defined as updating a single input neuron, necessitating N iterations to fully update all input neurons.

Considering *x* as our input, *W as* the weights matrix, and *t as* a chosen threshold value, a synchronous update follows the formula:

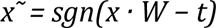

When performing an asynchronous update of the input *x* at randomly chosen position *j*, we compute the dot product between the input *x* and column *j* of the weights matrix. In other words, we considered all the neurons in position *j* in the patterns and their respective connectivity value. Thus, the formula is:

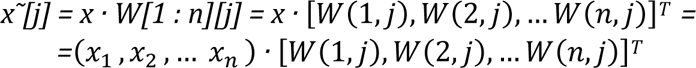

The output of this formula will yield a positive numerical value with a minimum value of 0. This occurs when no correlations are observed between the firing neurons in the input and the patterns associated with the specific position *j.* The resulting values increase proportionally to the strength of the correlation between the patterns, which directly influences the values in *W*. These values encapsulate the correlation between our input and the memorized patterns. Consequently, we applied a threshold to each value to convert it to 0 or 1.

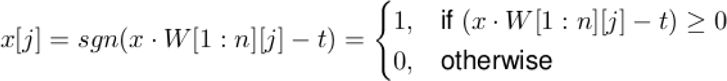

To create a memorized pattern, it is necessary to minimize to reach a stable state, hence reaching a stable state, which is the main idea behind the updating process of HNN (Koiran, 1994; Ramsauer et al., 2021; Šíma et al., 2000).

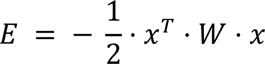

Each stored pattern has a corresponding minimum energy, which acts as an attractor. Therefore, it is logical to keep minimizing the energy function, as these are the states we aim to achieve with each update (Koiran, 1994; Ramsauer et al., 2021; Šíma et al., 2000). The weights matrix *W* comprises the connectivity values that indicate a noticeable tendency to increase while storing similar patterns. Therefore, the energy function also has minimums corresponding to spurious states - they are not patterns but modifications or linear combinations, which can potentially cause our input to become trapped (Koiran, 1994; Šíma et al., 2000).

## Implementation

The algorithmic implementation was conducted utilizing the Python programming language. All necessary features required for the network’s functionality were developed from the ground up, incorporating several libraries like numpy, matplotlib, and random, among others (Davison, 2008). The patterns denoted by ‘n’ were generated by randomly combining zeros and ones, resulting in a cumulative length of N neurons. The code availability is upon request.

### 1. Initial Experiments

The implementation process was initiated by constructing the system from its fundamental components, wherein various binary (i.e., 0s and 1s) and bipolar (i.e., 1s and -1s) patterns and inputs were explored through experimentation.

We selected the implementation of the binary HNN over the bipolar HNN because of its better alignment with the plausibility of neuron behavior, specifically in terms of firing when surpassing a specific activation threshold. The emphasis is placed on the magnitude of activation rather than the sign, as supported by previous studies (Litinskii, 2001; Weisbuch & Fogelman-Soulie, 1985). The preference for asynchronous neuron updates over synchronous updates is based on their advantage in facilitating the visualization of the network’s iterative process in shifting an input towards a stable state, as opposed to only perceiving the result in a single step (Aviel et al., 2004; Hopfield, 1982).

During this phase, the network would often attempt to update a specific neuron; this is not only redundant, as it would yield the same outcome, but also an insufficient depiction of how neurons activated during the recall process.

Initially, we ensured that each input neuron was updated at least once - all input positions were randomized at the beginning of a complete cycle of neuron updates and later used to update the input asynchronously (Fig.1b). Thus, a full input update after N neuron iterations was ensured. This asynchronous updating approach was subsequently employed in all subsequent implementations.

Afterward, we introduced perturbations of HNN’s memorized patterns to determine the expected outcome after a series of iterations. The algorithm stochastically selects a single pattern from the n patterns and perturbs its memory by flipping the state of some neurons, resulting in a change from 0 to 1 or vice versa. At first, the function was designed to accept a probability value representing the likelihood of each neuron being flipped. However, it was then determined that it would be more beneficial to specify the number of random neurons to be perturbed to gather data on the performance of the HNN. The patterns denoted as “n” were generated by randomly combining zeros and ones, resulting in a cumulative length of N neurons (Fig.1b).

After completing preliminary trials, we improved all the Python code and successfully developed a functional binary HNN. As a logical progression, we proceeded to gradually augment the number of patterns n and then analyzed the behavior of the HNN. It has been indicated that the memory capacity^1^ of the HNN decreases as the overlap of patterns increases (Hopfield, 1982; McEliece et al., 1987; Ramsauer et al., 2021). Hence, before implementing the threshold, we rechecked the computation of the weights matrix W, specifically how the outer products of the patterns expand.

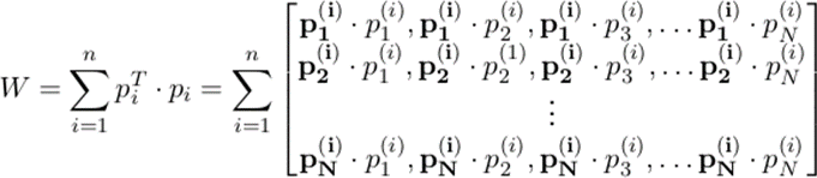

We applied the unfolded formulation of *W* to our asynchronous update of neuron j.

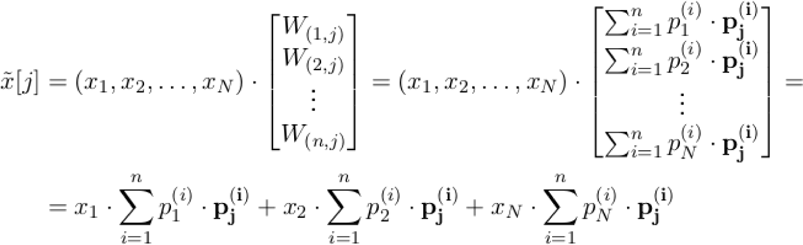

During this phase, it was observed that overlapping or correlated patterns had a detrimental effect on the recall rate. This was evident when multiple patterns yielded high values during a neuron update. For instance, in the scenario where 50% of the patterns exhibit neuron activity at position *j*, such noise would provide a significant challenge in retracing the original pattern from which perturbations were made.

Without any correlation between the patterns concerning the *j*-th neuron, the least achievable value is 0. This implies that if neurons are firing at position *j* in the patterns, the corresponding neurons in the input x are inactive, or no neurons are firing at location *j*.

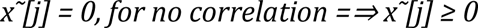

In the alternative scenario where all patterns exhibit activation in the *j*-th neuron, the resultant value can increase considerably, as it would be the total number of occurrences of the number one in each pattern. This would result in a strong correlation among all the stored concepts, as they share a common microfeature, specifically the firing of the *j*-th neuron. At the same time, the input *x* exhibits firing in all of its neurons. The purpose of analyzing this situation is simply to highlight the extensive spectrum of potential values for the updated returns; however, it is unlikely for this scenario to occur.

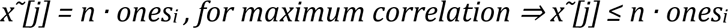

We consider *ones_i_* as the number of neurons firing in each pattern *p_i_*.

### 2. Pattern Suppression

We have an input that is a perturbation of pattern no. 1; this means that once we iterate our input through the network. Because our features are closest to those of the pattern we disturbed from, pattern no. 1, we anticipate returning to the stable state it offers.

To be creative and find another pattern to connect with, we need to ensure that our network will not let our input iterate towards pattern no. 1 while at the same time keeping it part of our system (Abe, 1993). In other words, we do not erase a concept or break its connections; instead, we shift our focus to finding another solution, and this is the rationale beyond reaching the idea of suppressing a pattern.

We can suppress a pattern from our network without deleting it by subtracting its contributions from the weight matrix W, which measures the correlation between patterns as perceived by each neuron. When performing a specific neuron update, we have to reduce the value of a given pattern to disable its contribution. Therefore, the new formula for updating neuron *j* of input *x* while suppressing pattern no. k, which will be later evaluated through a threshold, is:

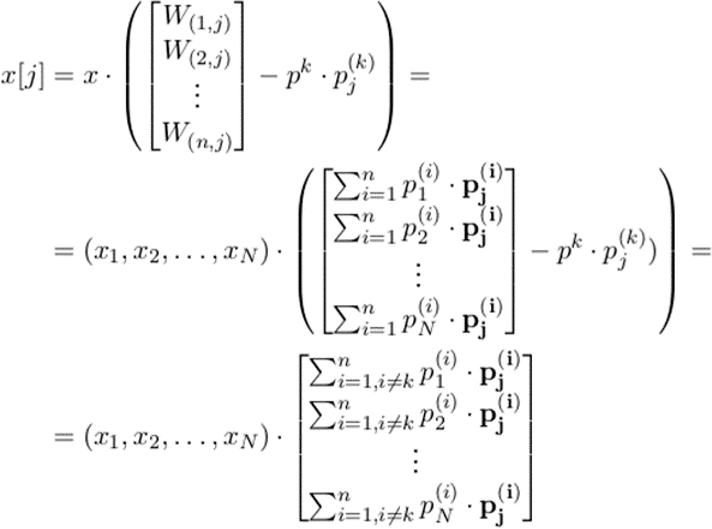

After this preliminary analysis, it can be inferred that the update operation yields a value that falls within, which increases significantly along with the number of patterns. As choosing an appropriate threshold that can discretize the result of the update computation is essential for the functionality of HNN, it is imperative to understand better the actual values to make informed decisions regarding threshold selection and neuron update. Here, we refer to the **“second Knob of creativity”** to make our network reach another solution. Twisting the knob suppressed specific patterns, allowing the HNN to creatively develop a pattern to associate our input with.

Throughout the implementation process, we realized the importance of the threshold in the binary HNN. Specifically, we observed that there was only a small window of range values for the threshold within which our network could attain a stable state while simultaneously suppressing certain patterns. One possible approach to address this issue would involve utilizing the first knob that was introduced and tweaking the threshold values gradually until our network exhibits creative behavior and ultimately converges into an alternative pattern.

### 3. Reconstructing Associative Chains

Semantic association-based creative thinking involves connecting previously unrelated notions to develop a new path (Beaty et al., 2023; Kenett & Beaty, 2023; Li et al., 2021; Luchini et al., 2023). For example, the input is a rose, and the output is honey. How can we find their associative chain? Our primary association is with a floral entity, specifically a rose, given its striking resemblance. Upon further contemplation of the various facets of flowers, we came to pollen as our next connection. We find the correlation between bees and pollen collection by considering what else we can relate to pollen, not flowers, roses, or being trapped pollen. We have yet to attain our objective, thus we explore what bees can connect to, except for roses, flowers, and pollen, and think about honey. These connections are still considered obvious and activated by slightly different groups of neurons, known as neural cliques (Wang et al., 2003).

The situation is different once our brain’s oblivious or already established connections do not solve our specific problem. Continuing the previous example, what if you have a bouquet of roses but no available vase - a new creative association could be to place them in a cup of water. When you are engaged in taking a step back from the situation and, as a result, make new connections, the group of neurons that fire together to form it are related to atypical microfeatures and are known in the neuroscience discipline as neurds (Gabora, 2010). Thus, another associative chain is created by combining one or more associative links, which tie together two concepts based on the neurons firing at common microfeatures.

## Results

### 1. Threshold Limits

We measured the success rate by counting how many tests resulted in a perfectly reconstructed memory. This network generates “n” new patterns with 10% of randomly positioned firing neurons, perturbs a randomly selected pattern with 100 neurons, and feeds it as input.

The threshold ranges with a success rate of 100, indicating that it effectively recalls the perturbed pattern and demonstrates the decreasing size of windows as n increases in all tests (Fig. 2). For values of n greater than or equal to 8, the network was not able to achieve a complete success rate. The success rate reduces dramatically for any threshold number. Specifically, for n = 8, the maximum success rate observed was 87%. For n = 9, the success rate dropped to 72%.

**Figure 2.**
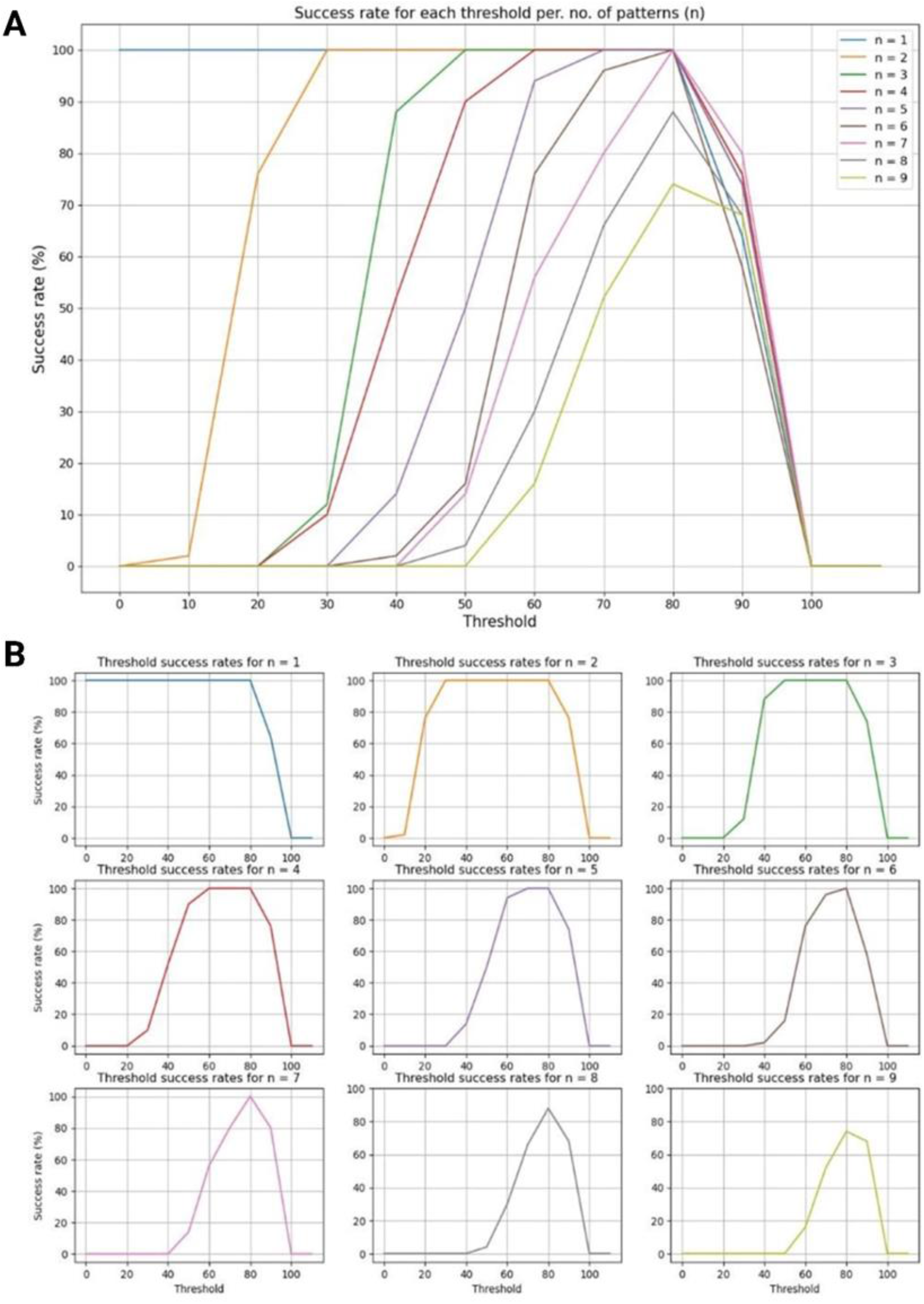
Illustration of measuring the success rate, quantifying the number of tests yielded in a perfectly reconstructed memory. Panel A illustrates the success rates of each threshold value for each number of patterns n. Panel B represents the success rates of each threshold value for each number of patterns n (i.e., the collected data in individual graphs, one for each value of n tested).

Remarkably, for all n for which a maximum success rate was achieved, the maximum threshold that would act is t = 80; the network completely shuts down for thresholds t ≥ 100, having a 0 recall rate. The threshold ranges of each *n,* which resulted in a perfect success rate of 100% or a good success rate (below 100 % but above 75% (Fig.3a), and the minimum and maximum values for which we achieve a complete success rate (Fig.3b).

**Figure 3.**
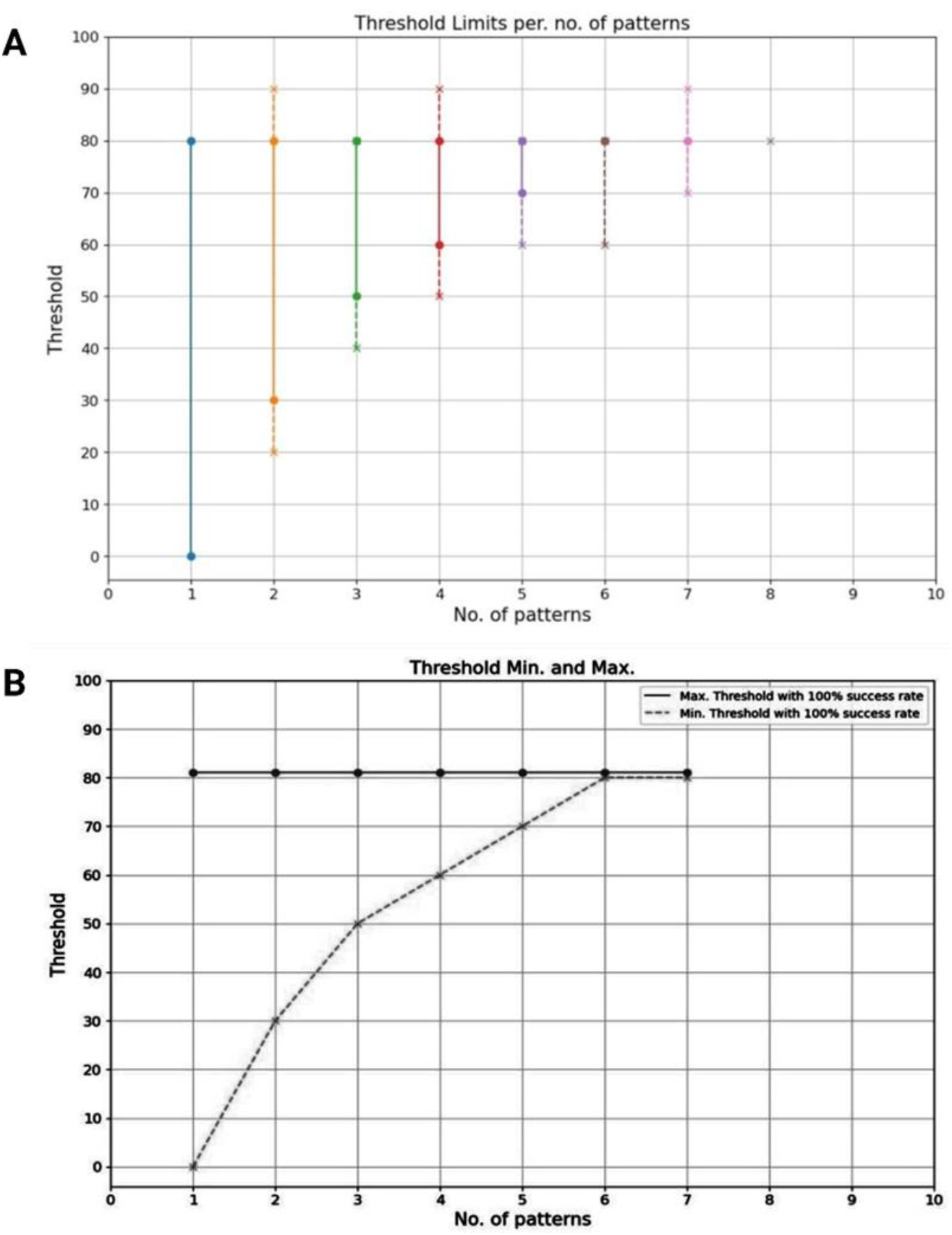
Depiction the successful threshold values. Panel A illustrates threshold ranges with a perfect success rate (=100%, solid lines) and reasonable success rate (≥75% and <100%, dotted lines). Panel B displays the lowest and highest threshold values with a reasonable success rate); it visualizes the almost logarithmic decrease of the ranges as n increases and the maximum threshold of t = 80 for a total success rate for all values of n.

This figure visualizes the almost logarithmic decrease of the ranges as n increases and the maximum threshold of *t* = 80 for a total success rate for all *n*.

The upper limit of the threshold can be elucidated by considering the 10% of firing neurons for each pattern and the perturbation of 100 neurons that we employ to generate our input. Thus, in each of our patterns, the probability of a neuron being 1 is 10%, while the probability of it being 0 is 90%. Therefore, if we flip 100 neurons, most likely 10% of them, which are ones, will become zeros; consequently, we have 10 neurons shutting down, and the rest of 90% of zeros will become ones; accordingly, we have 90 new firing neurons. This concludes that the input has approximately −10 + 90 = 80 new values of one.

When applying the thresholds, we attempt to shut down the extra neurons activated in the perturbation process to reach a stable state with approximately 80 fewer firing neurons. Therefore, it is logical to consider the maximum value obtained during the examination of the gathered data.

Finally, evaluating the gathered data underlines the network’s capacity; in particular, we can have a perfect recall rate for *n* ≤ 7. The deterioration of the reconstruction of patterns can be attributed to the cross-talk phenomenon, as discussed earlier. I.e., as the number of stored patterns increases, the likelihood of correlations between them also rises substantially. Consequently, retrieving a particular pattern that has been disrupted becomes more challenging.

### 2. First Knob of Creativity

We used the threshold as a parameter to twist this **“first knob of creativity”** to regulate whether the network can operate creatively, discover a solution, or shuts down completely. If we twist the knob to t = 60 for n = 3 memorized patterns, the network will act creatively and find a solution, but if we twist it to t = 35, it will shut off and not provide a solution. The network will act creatively for n = 5 patterns and find a solution if we twist it to t = 70, Fig.4.

**Figure 4.**
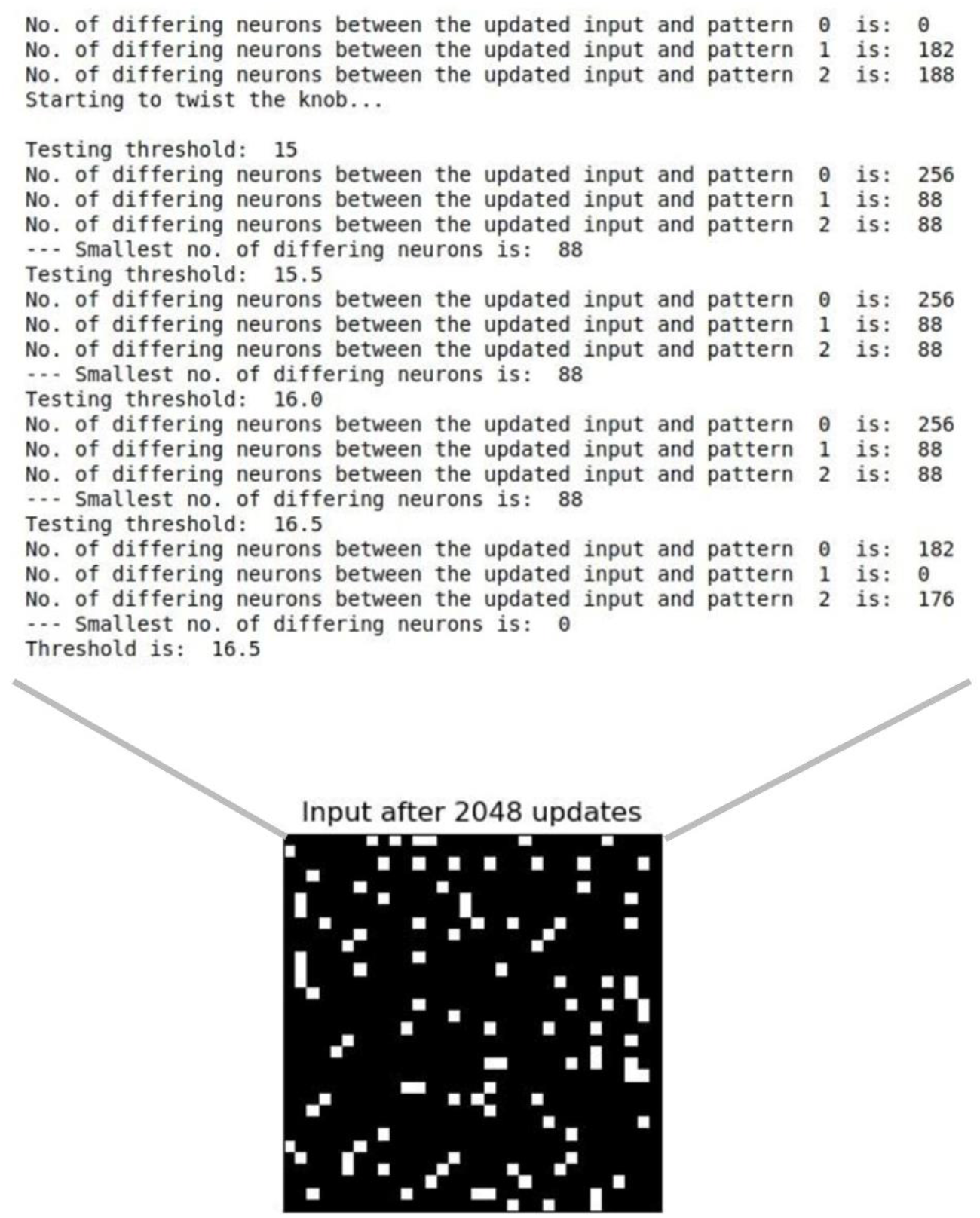
Illustratation if an instance where pattern suppression is effectively achieved. Initially, the input iterates towards pattern number 0, but upon suppressing this pattern by adjusting the first knob of creativity controlling the threshold, the input subsequently iterates towards pattern number 1.

### 3. The Second Knob of Creativity

We feed input into our network, which iterates toward its most associated pattern, in our case, pattern no. 6, establishing our first associative connection (Fig.5).

**Figure 5.**
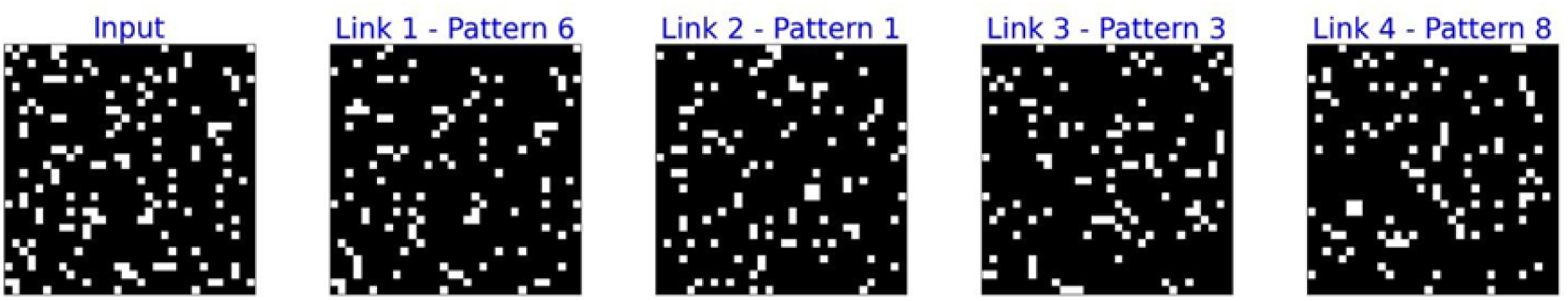
Depiction of the input pattern that exhibits the highest correlation, indicating distinct associative chains represented in various links. Pattern 6 is responsible for generating the initial associative link. We will input pattern number 6 into our network to locate the desired link while simultaneously inhibiting it. As result, HNN is compelled to establish an association between pattern number 6 and another. Consequently, the generation of the second link with pattern number 1 occurs. The validity of this new link is based on the assumption that there is a correlation between pattern number 6 and the initial input. Accordingly, the fundamental nature of the input is maintained during the process of identifying the future relationship. To extend the associative chain, we include pattern no. 1 as an input while excluding itself and pattern no. 6. This approach guarantees the identification of subsequent links, specifically with pattern no. 3, and so on. For a better understanding of these associations, see Figure 7.

We will suppress pattern 6 and feed it to our network to force the HNN to find a relationship between it and another pattern. The second link, with pattern no. 1, is generated. This new link is valid because pattern no. 6 and the initial input have correlated features; therefore, the input kernel is not lost when finding the next association. For a longer associative chain, supply pattern 1 as an input while suppressing itself and pattern 6, to locate the next link, especially with pattern 3, etc.

To advance our research, we evaluate our conceptual framework further using more patterns. However, the limited capacity of the binary HNN impeded our progress, prompting us to delve further into the modern HNN.

### 4. Modern Hopfield Network

The modern HNN was developed a few years ago, aiming to support continuous states implementing a new update rule and its respective energy function (Koiran, 1994; Krotov & Hopfield, 2016; Ramsauer et al., 2021; Šíma et al., 2000). Therefore, we introduced our perspective by rearranging the HNN into a two-layer network (Fig.6).

**Figure 6:**
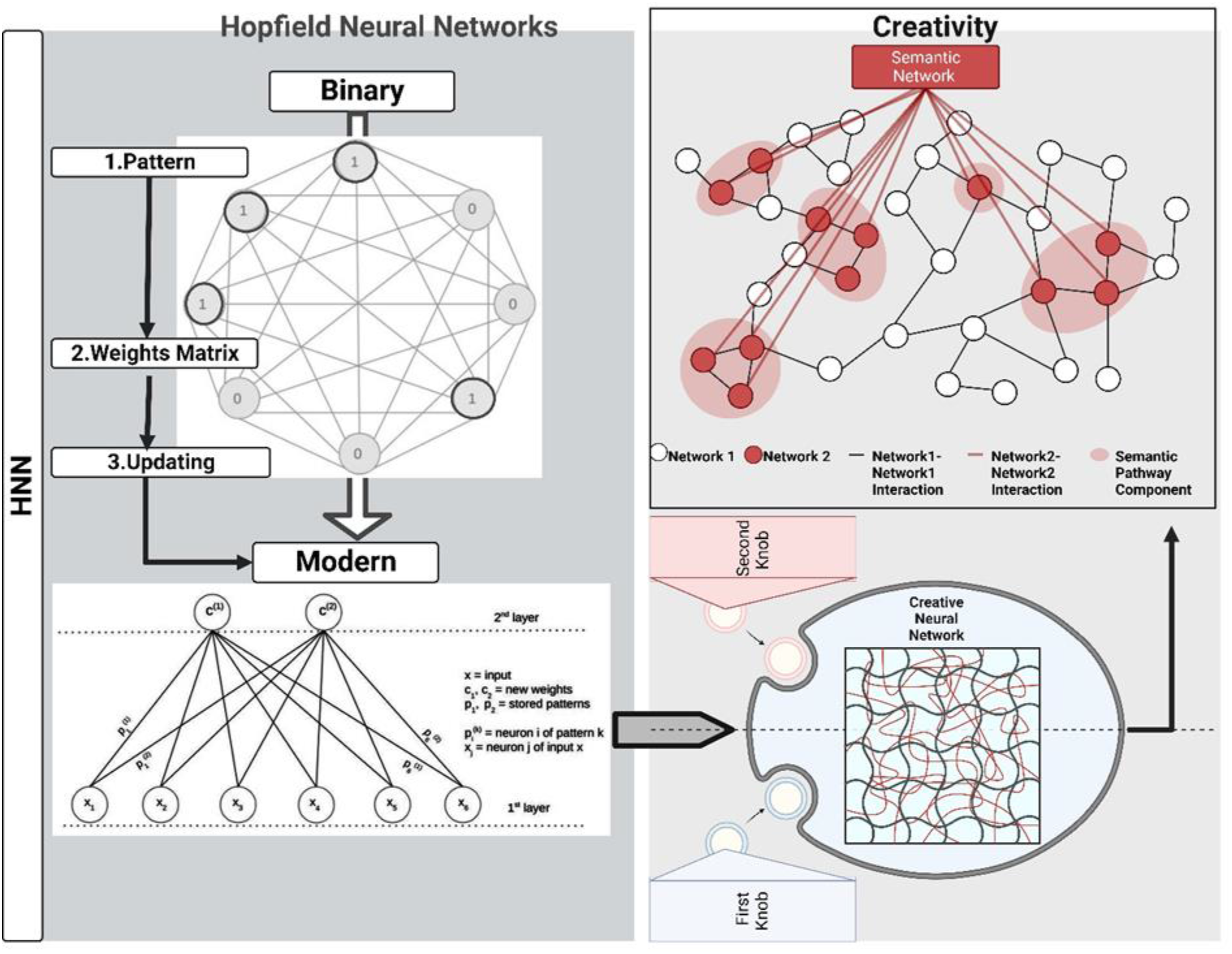
Summary of a hypothetical model of the experiment. Initially, we started with binary HNN, and as a result, we decided on modern HNN as a suitable model to represent the associative chains of creativity. We created modern HNN as a two-layer model. The first layer comprises the neurons of the input, and the connections between the two layers are realized through the neurons of the patterns. The top layer contains the core part of the updating process, which may be regarded as the “new weights,” denoted as c(i). The ‘new weights’ are a computation of the correlation between the current state of the input x and each pattern, later put through a normalization performed by the softmax function. Accordingly, this resembles the first and second Knobs of creativity, symbolized as a semantic association.

The first layer consists of input neurons, whereas the interconnections between the two layers are established through the neurons representing patterns. The second layer encompasses the fundamental aspect of the updating process, which we may presently regard as the “new weights,” denoted as c(i). These “new weights” are derived by calculating the correlation between the current state of the input variable x and each pattern. These weights are then normalized using the softmax function.

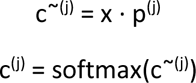

The softmax normalization function is defined as follows:

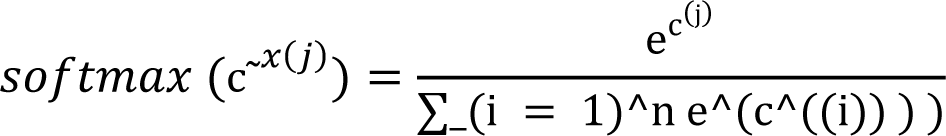

At the start of each asynchronous update, c was calculated to discover the most correlated pattern. The softmax function normalizes the pattern we should iterate; thus, setting an equal threshold is unnecessary. Accordingly, all current HNN implementations used t = 0.5.

We achieved a significantly higher recall rate and increased storage capacity using the modern HNN. We tested our network and got a perfect success rate even when storing 100 patterns. This first step is impressive compared to the simple HNN with 8 stored patterns as a capacity. The new updating rule improves performance using patterns’ neurons instead of the weights matrix. In modern HNNs, ‘new weights’ function as neurons, storing a single value from 0 to 1 due to normalization and dictating the idea activated, similar to a ‘decision’ neuron.

After successfully implementing the two HNN layers, we focused on associative chains. The most crucial part of their creation was suppressing a pattern in this new framework. We shut off ci = 0, the “decision” neuron in control of the thought we want to suppress, similar to memory reconstruction. Thus, we retained its linkages but focused on the other concepts, creating associative chains linking previously unrelated concepts in a framework with a higher capacity.

## Discussion and Concluding Remarks

We created associative connections between previously unconnected concepts using HNN to create a creativity-based semantic network conceptual framework (Fig.6). We started with simple binary and bipolar HNN and found the significance of the threshold; after performing a data analysis collection by testing several threshold values for different numbers of patterns, we defined the **“first knob of creativity”**, which controls whether the network acts creatively (i.e., finds a new path) or completely shuts down.

We also uncovered the capacity of the network n = 8, namely how many patterns we could store while still having a perfect recall rate. Here, we questioned what more we could do to help our system act creatively and reach another solution. At this stage, we defined suppressing a pattern such that it still exists in our binary HNN model and makes its connections. We stepped the theoretical step back and focused on finding another connection; thus, we established the **“second knob of creativity”** to assist our network in finding a solution other than suppression. Being constrained by the small capacity of the binary HNN, we transitioned to a modern HNN consisting of two layers. After redefining the suppression of a pattern in the new context, we successfully generated associations and tested our implementation using more patterns.

The first and second knobs assisted us in creating associative links between previously unrelated concepts and further developing them computationally into associative chains connecting several concepts; for a better understanding of the concept, see Fig.7.

**Figure 7.**
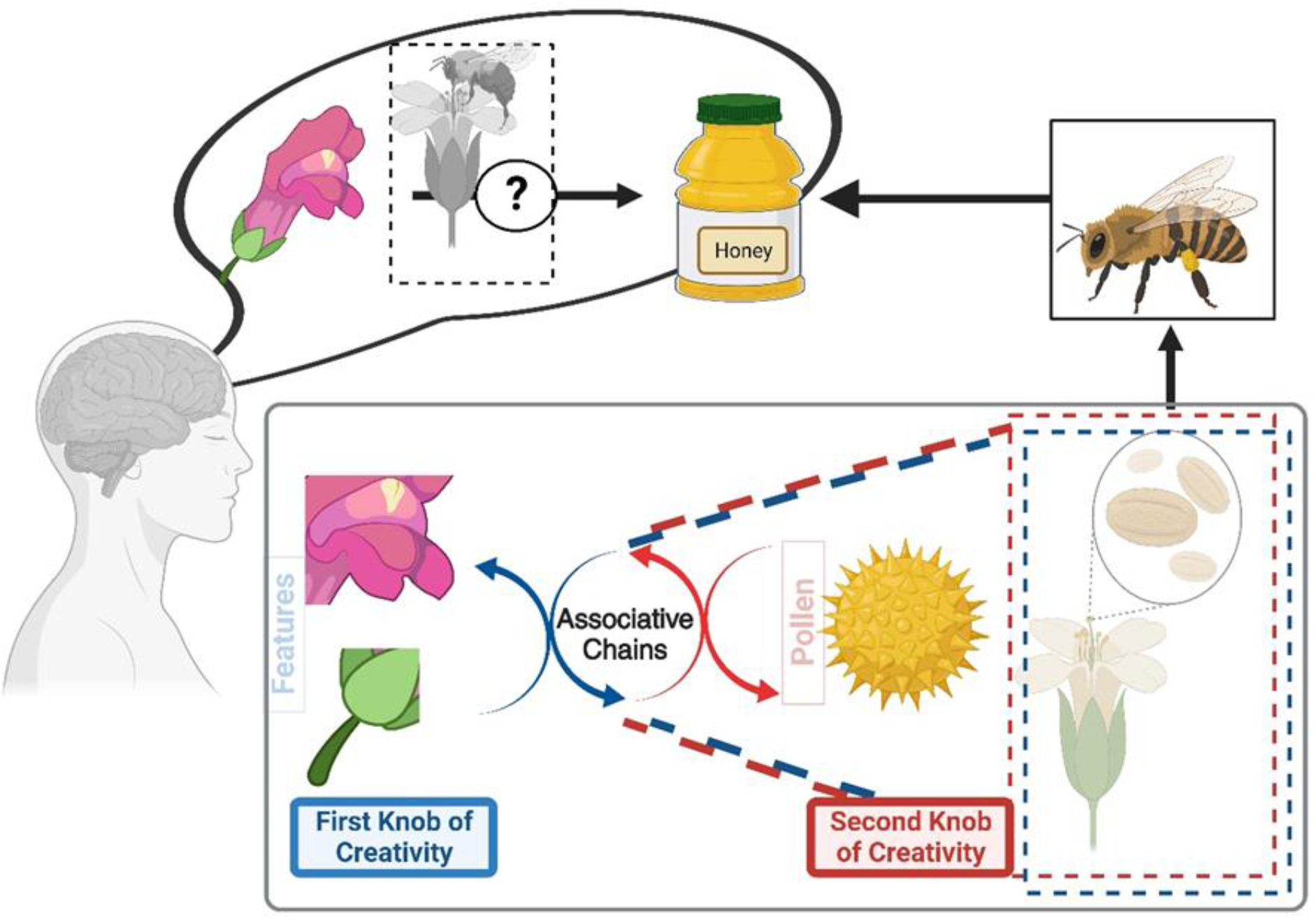
An illustrative case demonstrating the link of concepts and the potential extent of our associative chain without imposing a preconceived restriction on these links. If we were to examine the input “a rose” and the desired endpoint “honey,” what would be the associative chains? Which strategies can be utilized to determine the connection between these entities? The rose is classified as a flower and is mainly associated with the concept of flowers. Therefore, our initial connection is established with the category of flowers. In this context, the focus is directed towards the attributes of a flower, which may be considered the initial catalyst for **the first knob of creativity**. Pollen emerges as the subsequent element, representing the second facet of creative exploration (i.e., **the second knob of creativity**). In this context, we construct associative chains and engage in cognitive processes to identify additional associations with pollen while avoiding simplistic connections such as flowers, roses, or the pollen itself. Consequently, we discover a correlation between bees and their role in pollen collection. The desired objective has yet to be achieved, so further exploration is undertaken to ascertain the factors bees associate with accessing honey, including the bee, flower, and pollen.

In conclusion, we successfully presented for the first time a neurocomputational framework model for creativity-based semantic associations, being implemented in both a simple, binary HNN and a modern, two-layer one (Fig.6 and Fig.7). A potential explanation of our findings is that our neurons take a step back from the contextual focus and find alternatives associated with seemingly unrelated concepts (Beaty & Kenett, 2023; Gabora, 2010; Krotov & Hopfield, 2016; McEliece et al., 1987; Mednick, 1962) when analytical thinking is insufficient.

Accordingly, we do not have to search the whole network and compare our input with all the patterns to find a solution; it helps us retrieve the memorized items seamlessly. Therefore, an associative link is created when a group of neurons (i.e., neural cliques; Wang et al., 2003) fire up together while being part of both concepts we are connecting; this allows us to find a solution that would not be an oblivious choice but that solves our problem This could be analog of creative thinking based on semantic associations, allowing us to search on the lower levels of memories to find which concepts relate even on a micro-level to our situation (Beaty et al., 2023; Kenett & Beaty, 2023; Li et al., 2021; Luchini et al., 2023). The patterns and how we store and reconstruct them align with the biological processes, as human memory is also distributed, coarse-coded, and content-addressable (Beaty et al., 2023; Benedek et al., 2023; Gerver et al., 2023; McEliece et al., 1987).

This preliminary computational neural network model has numerous possibilities for investigation, as we can find creative ways to connect concepts and see how far our chain can reach without imposing a predetermined limit on the extent of these connections.

## Future Direction

Several interesting open questions need to be addressed in the upcoming research work. For instance, it would be ideal for testing the framework on patterns representative of real-life concepts (i.e., create a dataset for them and define a set of multiple features that should be defined for each of them (e.g., human/animal/object, weight, height, color, etc.)) as a next step, daily creativity (Richards, 1993; Runco & Bahleda, 1986); however, it is challenging empirically. However, it is possible through implementing an algorithm that takes as input a concept, and after the user answers questions about its features, we can collect the data and transform it into a pattern. We understand the idea of a feature as a group of neurons (e.g., the first 10 neurons of each pattern) and the definition of it as the positions of the firing neurons (e.g., a neuron firing on the second position in a weight feature translates to a light object, while 0 means a weight is not applicable).

Therefore, a natural and crucial next step would be testing this theoretical framework on thoughts modeled by real-life concepts; accordingly, the network’s creativity-based semantic associations will also be easier to observe. Taking it a step further, we could also redefine the way associative links get created by iterating solely on the defined features and, as such, suppressing certain features.

## Footnotes

1 We refer to the memory capacity as the maximum value for the number of patterns n, for which the inputs can reach a stable state, whereas the correlation between the patterns causes noise (Krotov & Hopfield, 2016).

